# *Barley stripe mosaic virus* γb protein targets thioredoxin h-type 1 to dampen SA-mediated antiviral defenses

**DOI:** 10.1101/2022.01.06.475245

**Authors:** Zhihao Jiang, Xuejiao Jin, Meng Yang, Qinglin Pi, Qing Cao, Zhenggang Li, Yongliang Zhang, Xian-Bing Wang, Chenggui Han, Jialin Yu, Dawei Li

## Abstract

Salicylic acid (SA) acts as a signaling molecule to perceive and defend against pathogen infections. Accordingly, pathogens evolve versatile strategies to disrupt the SA-mediated signal transduction. However, it is necessary to further characterize how plant viruses manipulate the SA-dependent defense responses. Here, we show that *Barley stripe mosaic virus* (BSMV) infection activates SA-mediated defense signaling pathway and upregulates the expression of *Nicotiana benthamiana* thioredoxin h-type 1 (NbTRXh1). The γb protein interacts directly with NbTRXh1 *in vivo* and *in vitro*. Overexpression of NbTRXh1, but not a reductase-defective mutant, impedes BSMV infection, whereas low *NbTRXh1* expression level results in increased viral accumulation. Similar with its orthologues in *Arabidopsis*, NbTRXh1 also plays an essential role in SA signaling transduction in *N. benthamiana*. To counteract NbTRXh1-mediated defenses, the BSMV γb protein targets NbTRXh1 to dampen its reductase activity and thereby impairing downstream SA defense genes expression to optimize viral cell-to-cell movement. We also found that NbTRXh1-mediated resistance defends against *Lychnis ringspot virus*, *Beet black scorch virus*, and *Beet necrotic yellow vein virus*. Taken together, our results reveal a novel role for the multifunctional γb protein in counteracting plant defense responses, and broadens the broad-spectrum antibiotic role of SA signaling pathway.

**One sentence summary:** BSMV γb protein impairs NbTRXh1 reductase activity and dampen downstream SA-related genes expression to facilitate viral cell-to-cell movement.

## Introduction

Salicylic acid (SA) is a key hormone for plants to defend against biotrophic and semi-biotrophic pathogens (Ding and Ding, 2020; Zhao and Li, 2021). Upon pathogen invasion, redox status in plant cell changes and activates thioredoxins (TRXs). Consequently, NONEXPRESSOR OF PATHOGENESIS-RELATED GENE 1 (NPR1), a master regulator of SA-mediated defense pathway (Cao et al., 1997), is translocated to nucleus where activates downstream defense genes expression in *Arabidopsis* (Mou et al., 2003). Besides its role as transcriptional coactivator, NPR1 is also involved in a series of defense events such as callose deposition, reactive oxygen species (ROS) production, and systemic acquired resistance (SAR) activation (Peng et al., 2021).

TRXs are ubiquitous, low-molecular-mass proteins with two cysteines in their conserved active site WC(X)PC which are involved in numerous biochemical processes in cells and widely distributed in plants, animals, bacteria and yeasts (Mata-Pérez and Spoel, 2019; Kumari et al., 2021). The subcellular localization of TRXs isoforms varies greatly in organelles. Thioredoxin h-type proteins (TRXhs), which constitute the largest TRXs subfamily, are primarily distributed in the cytoplasm (Kang et al., 2019). In *Arabidopsis*, AtTRXh3 and AtTRXh5 are required for catalyzing the conversion of NPR1 oligomers to monomers and subsequent NPR1 translocation to nucleus (Tada et al., 2008). Besides the investigation on their roles in reducing the oxidized thiols in NPR1, AtTRXh5 also acts as a selective protein-SNO reductase and regulates SA-responsive genes expression in plant immunity (Kneeshaw et al., 2014). Importantly, NPR1 from tabacco differs from AtNPR1 on lacking the conserved residues for cytoplasmic oligomer-nuclear monomer exchange and SA-dependent transcriptional activation (Maier et al., 2011); thus, whether TRXhs in tobacco function in SA signaling pathway is still unknown.

Given the essential role of SA in plant immunity, many pathogens-derived effectors have been identified to dampen SA-mediated immunity response via different strategies (Lorang et al., 2012; Mukaihara et al., 2016; Chen et al., 2017; Qi et al., 2018; Ji et al., 2020; Medina-Puche et al., 2020). In terms of plant viruses, the C4 protein from the geminivirus *Tomato yellow leaf curl virus* (TYLCV) re-localizes from the plasma membrane to chloroplasts to disturb SA biosynthesis upon activation of defense (Medina-Puche et al., 2020). *Tobacco mosaic virus* (TMV) replicase interacts with NAC domain transcription factor ATAF2 to suppress SA-related genes expression as a means to promote systemic virus infection (Wang et al., 2009). Also, SUMOylation of *Turnip mosaic virus* (TuMV) NIb protein suppresses the NPR1-mediated immune response via interactions with small ubiquitin-like modifier3 (SUMO3) (Cheng et al., 2017). Although some progress has been made revealing the battle between plant viruses and SA defense pathway, more in-depth research remains to be done.

*Barley stripe mosaic virus* (BSMV) is a positive-strand RNA virus containing three genome RNA segments designated RNAα, RNAβ, and RNAγ. RNAα and RNAγ are sufficient for viral replication in plants; RNAβ is required for virion assembly and movement process (Jiang et al., 2021). Recent advances give us a more complete understanding of the versatile roles of the γb protein in viral pathogenesis and the molecular interactions between the virus and the host. BSMV γb interacts with viral replicase αa to promote BSMV replication (Zhang et al., 2017), and interacts with movement protein TGB1 to enhance viral cell-to-cell movement (Jiang et al., 2020). In addition, BSMV γb protein subverts autophagy-mediated antiviral defenses by impeding the interaction between autophagy key factors ATG7 and ATG8 (Yang et al., 2018), suppresses the peroxisomal-derived ROS bursts by interacting with glycolate oxidase (Yang et al., 2018), inhibits host RNA silencing and cell death responses through PKA-like kinase-mediated phosphorylation (Zhang et al., 2018), and disrupts chloroplast antioxidant defenses by targeting NADPH-dependent thioredoxin reductase C (NTRC) (Wang et al., 2021) to facilitate virus infection. Intriguingly, we also found plant counter-counter-defense strategy as serine/threonine/tyrosine kinase STY46 interacts with and phosphorylates γb to defend against BSMV infections (Zhang et al., 2021).

In this study, we revealed that BSMV infection interferes with SA antiviral defenses pathway. The γb protein targets NbTRXh1 protein and suppresses SA-mediated plant resistance by weakening the NbTRXh1 reductase activity, thereby inhibiting downstream defense genes expression, such as *pathogenesis-related gene 1* (*PR1*) and *PR2*, to optimize BSMV cell-to-cell movement. We also found that NbTRXh1 plays a general antiviral role against plant viruses within different genera. Our results reveal a novel role of the multifaceted γb protein in SA-mediated antiviral responses to BSMV infection, and these findings significantly deepen our understanding of co-evolutionary arms race between plant defenses and viral counter defense.

## Results

### γb protein counteracts BSMV-induced SA defense responses

We previously found that SA signaling pathway respond to BSMV infections (Zhang et al., 2018). To systematically investigate the role of the SA-mediated defense pathway in BSMV infection, we measured the transcript level of SA-related marker genes during BSMV infections. These results showed that *NPR1* (Cao et al., 1994), *WRKY70* (Li et al., 2004), and *PR1* (Alexander et al., 1993) transcripts were significantly upregulated in BSMV-infected *Nicotiana benthamiana* plants compared with those in mock-inoculated plants (Figure 1A). Moreover, 0.5 mM SA treatment dramatically decreases BSMV proteins accumulation (Figure 1B). These results suggest that BSMV infection activates SA antiviral defense pathway.

**Figure 1.**
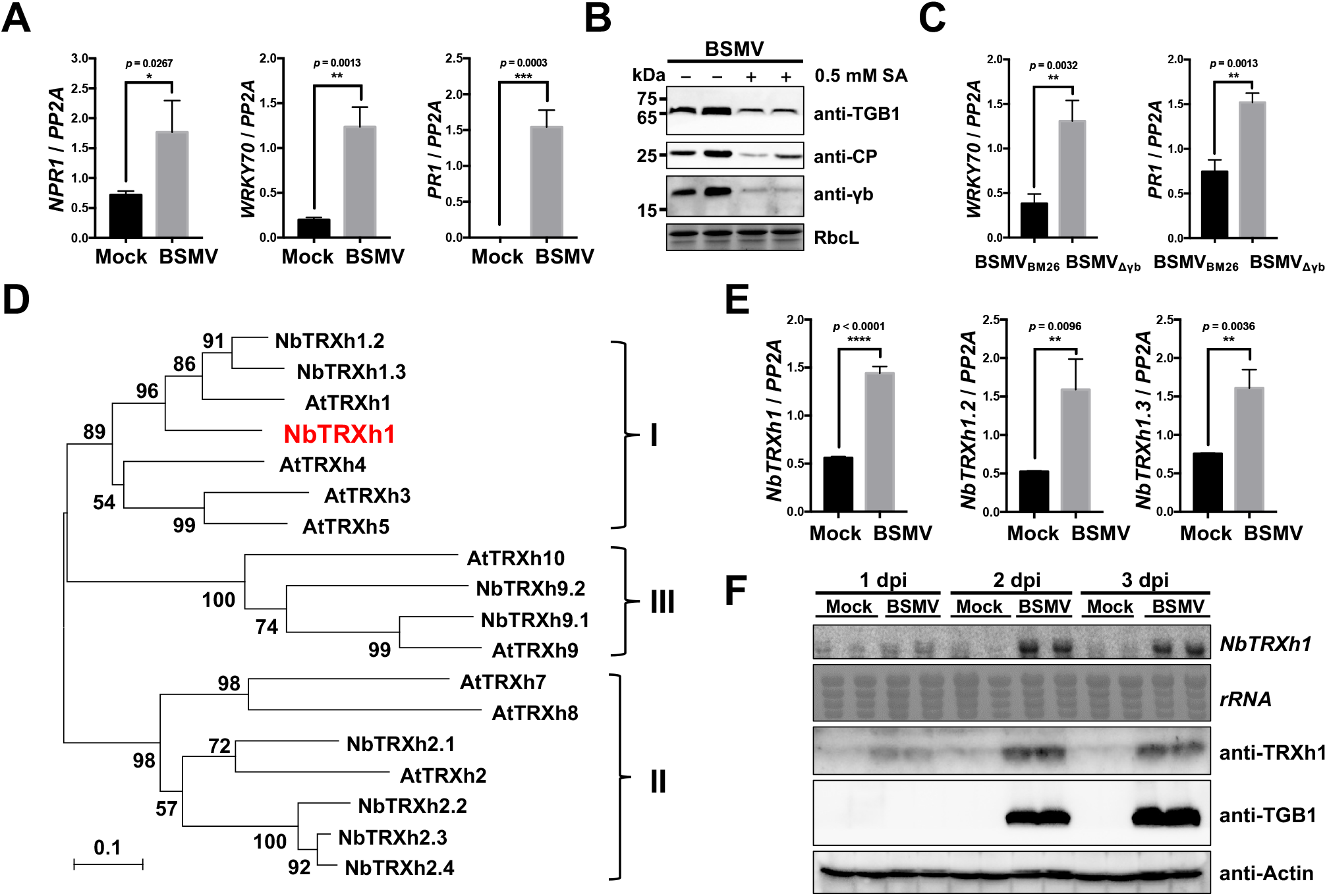
BSMV γb suppresses the SA signaling pathway during viral infection. **(A)** RT-qPCR analysis of the transcript levels of SA pathway-related genes in mock-inoculated or BSMV-infected *N. benthamiana* plants. The *protein phosphatase 2A* (*PP2A*) gene was used as internal control. Data were analyzed by Student’s *t* test (*, *p* < 0.05; **, *p* < 0.01; ***, *p* < 0.001). **(B)** Immunoblot analysis of BSMV-encoded proteins accumulation in *N. benthamiana* with or without 0.5 mM SA treatment. RbcL served as loading control. **(C)** RT-qPCR analysis of the transcript levels of SA pathway-related genes in BSMV_BM26_- or BSMV_Δγb_-infected *N. benthamiana* plants. Data were analyzed by Student’s *t* test (**, *p* < 0.01). **(D)** Phylogenetic analysis of TRXh proteins from *N. benthamiana* (NbTRXh) and *A. thaliana* (AtTRXh). The phylogenetic tree was constructed using neighbor-joining method, with the bootstrap analyses of 1000 cycles. The TRXh proteins were divided into three major subgroups I, II, and III as indicated. Bar represents sequence divergence of 0.1 amino acids. **(E)** RT-qPCR analysis of the transcript levels of *NbTRXh1*, *NbTRXh1.2*, and *NbTRXh1.3* in mock-inoculated or BSMV-infected *N. benthamiana* plants. Data were analyzed by Student’s *t* test (**, *p* < 0.01, ****, *p* < 0.0001). **(F)** Time course analysis of the transcript level and protein accumulation of *NbTRXh1* in mock-inoculated or BSMV-infected *N. benthamiana* plants by northern blot assay at 1 dpi, 2 dpi, and 3 dpi, respectively. Actin was used as loading control.

Since γb protein is a determinant of BSMV pathogenesis (Donald and Jackson, 1994; Bragg et al., 2004), we posited that the activation of SA pathways maybe associated with γb protein. However, owing to the significant decreased viral accumulation of BSMV_Δγb_ mutant (remove the whole γb ORF) (Zhang et al., 2017), compared with the wild type virus (Supplemental Figure S1), the comparison of the two viruses cannot correctly reflect the effect of γb on SA stimulation. Instead, we used a BSMV_BM26_ mutant of which the viral accumulation is similar to that of BSMV_Δγb_ (Supplemental Figure S1). BSMV_BM26_ mutant contains two basic motif mutations within γb (^25^RK^26^ → ^25^QN^26^) that abolish the RNA binding activity (Donald and Jackson, 1996). BSMV_BM26_ or BSMV_Δγb_ were inoculated into *N. benthamiana* by agroinfiltration and SA marker genes were analyzed by quantitative real-time PCR (RT-qPCR) at 3 dpi. The results showed that γb can suppress the SA defense pathway as verified by the remarkable downregulation of the transcript levels of *WRKY70* and *PR1* in BSMV_BM26_-infected *N. benthamiana* plants (Figure 1C). Altogether, these data indicate an additional role of γb in inhibition of virus-triggered SA defense responses.

### BSMV infection upregulates NbTRXh1 expression

To investigate how γb disturbs SA defense pathway, we performed a yeast two-hybrid (Y2H) screen using BSMV γb protein as a bait against a mixed cDNA library from tobacco plants as described previously (Zhang et al., 2021). *N. benthamiana* thioredoxin h-type 1 (NbTRXh1, GenBank accession: GQ354821) was identified as a potential γb-interactor. The full-length cDNA of NbTRXh1 was cloned based on the updated *N. benthamiana* genome database (https://ora.ox.ac.uk/objects/uuid:f34c90af-9a2a-4279-a6d2-09cbdcb323a2) (Kourelis et al., 2019), and the amino acid sequence of TRXhs from *N. benthamiana* and *A. thaliana* were analyzed. These proteins shared a conserved motif WC(X)PC, while contained variable N- and C-terminus, suggesting that these proteins might execute versatile functions in plant cells (Supplemental Figure S2). Intriguingly, nine putative NbTRXh paralogs were identified from updated *N. benthamiana* genome dataset (Kourelis et al., 2019) (Table 1). Subsequently, we generated a phylogenetic tree including the *N. benthamiana* NbTRXhs and *Arabidopsis* AtTRXhs assigning corresponding names for NbTRXhs based on the AtTRXhs orthologs. The NbTRXh1 showed high homology with AtTRXh1, AtTRXh3, and AtTRXh5, all of which are clustered into subgroup I (Figure 1D). A previous study has demonstrated that AtTRXh3 and AtTRXh5 catalyze SA-induced NPR1 oligomer-to-monomer switch (Tada et al., 2008), alluding to the functional link of NbTRXh1 with SA defense pathway in *N. benthamiana*.

**Table 1.** General information of NbTRXhs.

| Gene name | Gene ID<sup>a</sup> | Type | CDS (bp) |
| --- | --- | --- | --- |
| NbTRXh1 | NbD046153.1 | I | 369 |
| NbTRXh1.2 | NbD034443.1 | I | 354 |
| NbTRXh1.3 | NbD029648.1 | I | 351 |
| NbTRXh2.1 | NbD052469.1 | II | 426 |
| NbTRXh2.2 | NbD015627.1 | II | 390<sup>b</sup> |
| NbTRXh2.3 | NbE03054613.1 | II | 408 |
| NbTRXh2.4 | NbD015628.1 | II | 420 |
| NbTRXh9.1 | NbD027853.1 | III | 456 |
| NbTRXh9.2 | NbD031089.1 | III | 417 |

<sup>a</sup> The source where these IDs were taken, <sup>b</sup> Partial sequence.

Subsequently, to elucidate whether BSMV infection upregulates the expression of *NbTRXhs* in subgroup I, the transcript levels of *NbTRXh1*, *NbTRXh1.2*, and *NbTRXh1.3* were tested by RT-qPCR analyses. All the three genes were significantly upregulated in the context of BSMV infection (Figure 1E). Since the NbTRXh1 shares a high similarity with NbTRXh1.2 and NbTRXh1.3 (89.43%), we focused our further investigation on *NbTRXh1*. To elucidate the dynamics of the expression pattern of NbTRXh1 during BSMV infection, NbTRXh1 mRNA and protein levels were analyzed at 1-day post-inoculation (dpi), 2 dpi, and 3 dpi, respectively. The results showed that the expression of NbTRXh1 was induced as early as 1 dpi, reaching a maximun at 2 dpi, and maintaining a high level at 3 dpi (Figure 1F), suggesting that NbTRXh1 could respond to virus infection at an early stage.

### BSMV γb interacts physically with NbTRXh1

To confirm the results from the previous screen, Y2H assays were performed to test the interaction between γb and NbTRXh1. Despite the wild-type version of the NbTRXh1 protein failed to interact with γb, we found that a truncated mutant NbTRXh148-80, lacking for catalytic center, was sufficient to bind to γb (Figure 2A). To clarify whether the catalytic center (^42^WCGPC^46^) of NbTRXh1 is required for its interaction with γb, NbTRXh1-3xFlag or its enzymatic-defective mutant NbTRXh1_SS_-3xFlag (^42^WCGPC^46^ → ^42^WSGPS^46^) were expressed in BSMV_γb-GFP_ or BSMV_Δγb-GFP_-infected *N. benthamiana* plants (Zhang et al., 2017). The co-immunoprecipitation (co-IP) result showed that the γb-GFP co-precipitated with both NbTRXh1 and NbTRXh1_SS_ proteins, but not with GFP alone (Figure 2B).

**Figure 2.**
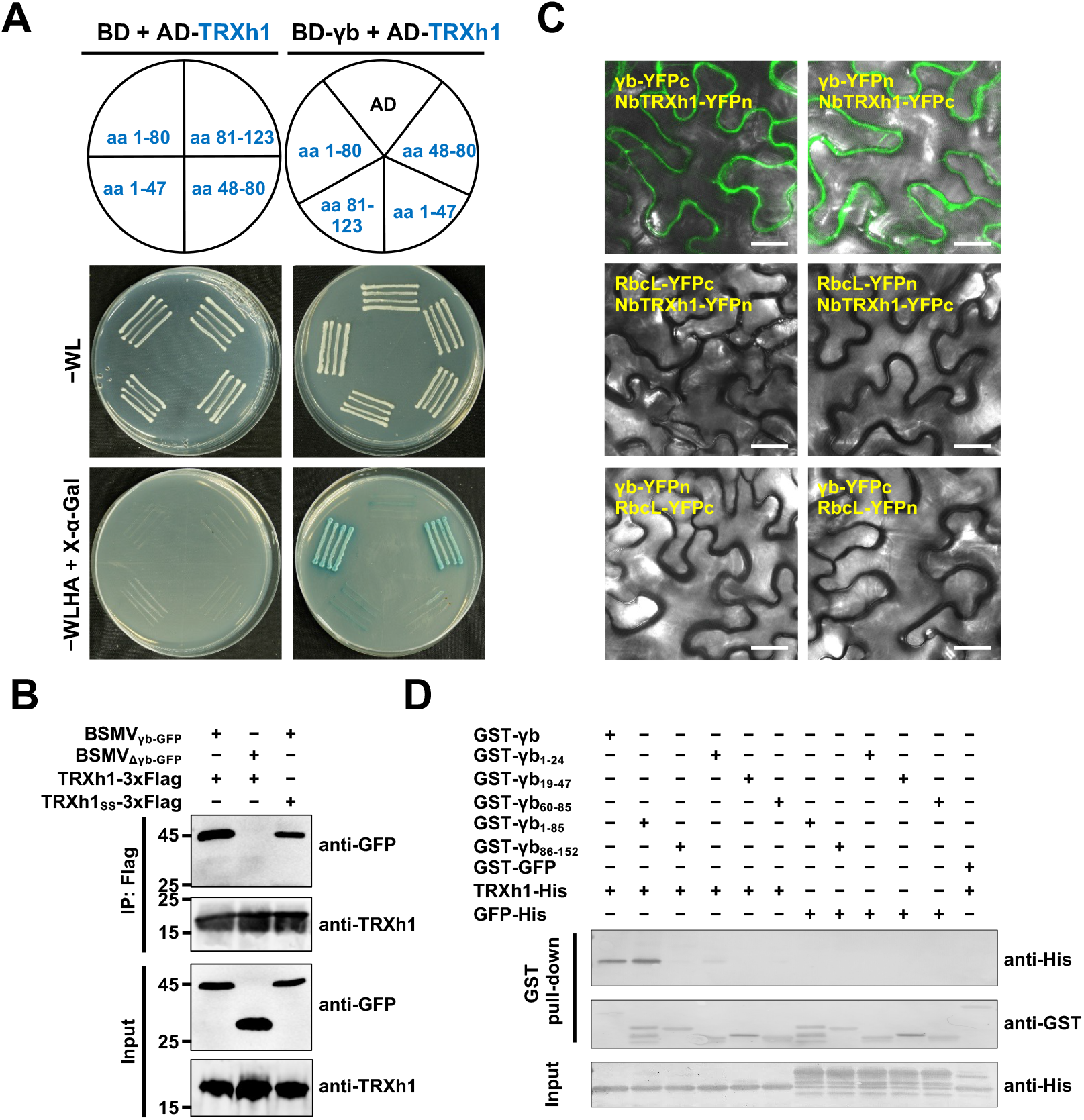
BSMV γb interacts with NbTRXh1. **(A)** Yeast two-hybrid assay to investigate the interaction of γb with NbTRXh1 and its truncated mutant. **(B)** Co-immunoprecipitation analysis to confirm the interaction between γb and NbTRXh1. *N. benthamiana* leaves infiltrated with different constructs were collected at 3 dpi. Total protein was immunoprecipitated with anti-Flag beads and analyzed by western blot with anti-GFP and anti-TRXh1 antibodies. **(C)** Bimolecular fluorescent complementation assay to test the *in vivo* interaction between γb and NbTRXh1 in *N. benthamiana*. The reconstructed YFP signal was visualized by confocal microscope at 3 dpi and depicted as a false-green color. RbcL-YFPn and RbcL-YFPc were used as negative controls. Bars, 20 μm. **(D)** GST pull-down assay to analyze the *in vitro* γb and NbTRXh1 interaction. Purified GST-γb or a series of γb truncated proteins were incubated with NbTRXh1-His or GFP-His. After being immunoprecipitated with glutathione-sepharose beads, the proteins were detected by western blot with anti-His and anti-GST antibodies.

Next, bimolecular fluorescence complementation (BiFC) assay was performed and the results showed that reconstructed YFP fluorescence signal appeared in *N. benthamiana* leaves co-infiltrated with γb-YFPc/NbTRXh1-YFPn and γb-YFPn/NbTRXh1-YFPc pairs, whereas the RuBisCo large subunit (RbcL, negative control) failed to interact with neither γb nor NbTRXh1 (Figure 2C). Protein accumulation was confirmed by western blot analysis (Supplemental Figure S3). Furthermore, we found that BSMV γb has no effect on the nucleo-cytoplasmic subcellular localization pattern of NbTRXh1 (Supplemental Figure S4).

To investigate the physical interaction between γb and NbTRXh1 *in vitro*, GST pull-down assay was carried out by using recombinant proteins purified from *E. coli*. The results showed that NbTRXh1 was specifically pulled down by GST-γb and GST-γb_1-85_ truncated derivative, and a faint shadow was seen in γb_1-24_ truncation mutant lane, whereas other truncations of γb did not (Figure 2D). In addition, we further indicated that γb_BM26_ protein, a crucial control protein in this study, can also interact with NbTRXh1 in GST pull-down assay (Supplemental Figure S5). Collectively, these data suggest that γb physically interacts with NbTRXh1 in cytoplasm and the catalytic center of NbTRXh1 is not required for the interaction.

### NbTRXh1 inhibits BSMV local and systemic infection in *N. benthamiana*

To understand the function of NbTRXh1 during BSMV infection, *N. benthamiana* leaves were co-infiltrated with *Agrobacterium* harboring a BSMV_γb-GFP_ infectious clone and NbTRXh1-3xFlag or NbTRXh1_SS_-3xFlag (Figure 3A). At 3 dpi, the proportion of GFP-positive cells in the regions expressing NbTRXh1-3xFlag was significantly lower than that in the regions expressing NbTRXh1_SS_-3xFlag (Figure 3B and 3C). Furthermore, viral proteins and RNA accumulations were significantly decreased in plants expressing NbTRXh1-3xFlag compared with those expressing NbTRXh1_SS_-3xFlag or mock-inoculated plants (Figure 3D), suggesting that the reductase activity of NbTRXh1 is essential to inhibit BSMV infection.

**Figure 3.**
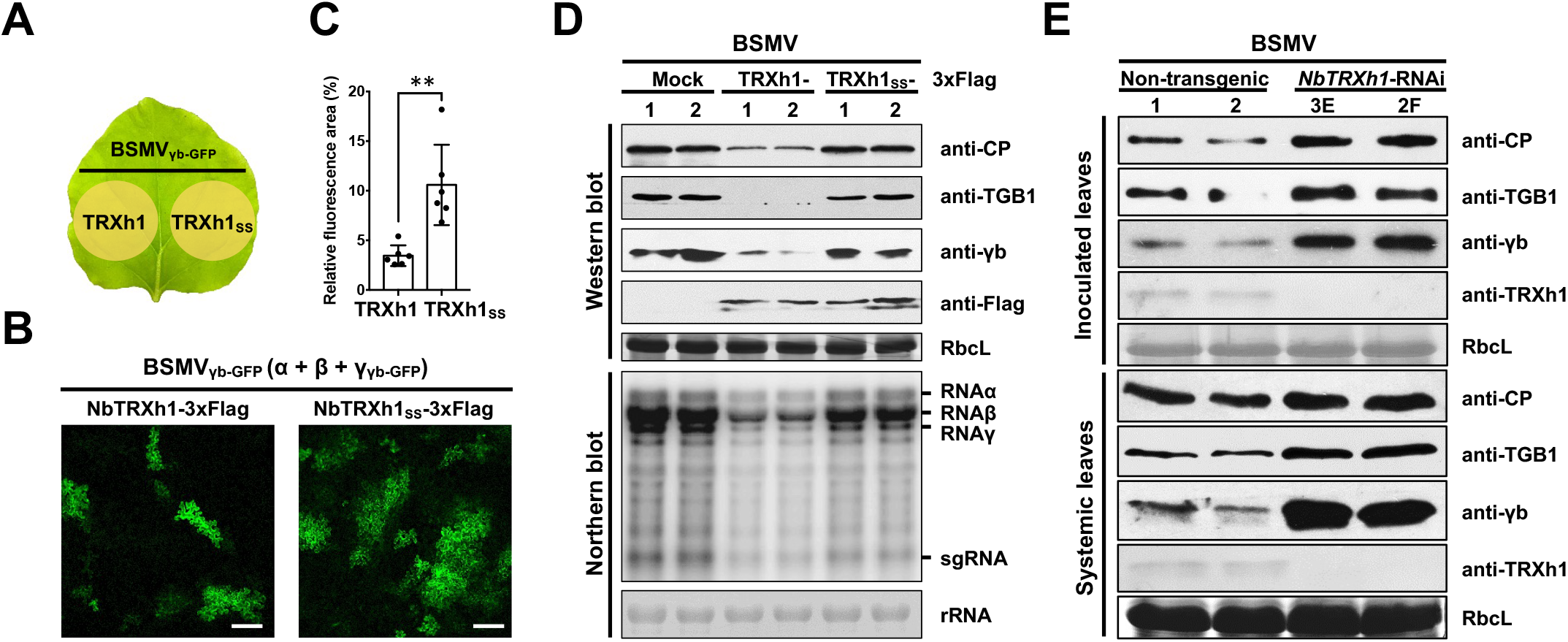
NbTRXh1 reductase activity reduces BSMV accumulation in *N. benthamiana*. **(A)** Schematic diagram of the half-leaf method used in Figure 3B. **(B)** Effects of NbTRXh1-overexpression on BSMV infection. BSMV_γb-GFP_ was agroinfiltrated into *N. benthamiana* together with TRXh1-3xFlag or TRXh1_SS_-3xFlag reductase-defective mutant. The GFP fluorescence indicates BSMV-infected cells and was photographed by confocal microscope at 2.5 dpi. Bars, 20 μm. **(C)** Quantification of the GFP fluorescent area in Figure 3B by using ImageJ. Data were statistically analyzed by Student’s *t* test (n = 6, **, *p* < 0.01). **(D)** BSMV accumulation was detected by western and northern blotting in *N. benthamiana* leaves agroinfiltrated with empty vector (EV), NbTRXh1-3xFlag, or NbTRXh1_SS_-3xFlag. Antibodies were shown on the right of the panel. Experiments were repeated twice. **(E)** Western blot showing BSMV accumulation in non-transgenic or *NbTRXh1*-knockdown *N. benthamiana* plants. BSMV was agroinoculated. Local and systemic infected leaves were collected at 2 dpi and 6 dpi, respectively. Antibodies were shown on the right of the panel. Experiments were repeated three times.

To confirm the anti-BSMV role of NbTRXh1, a short-hairpin RNA-mediated RNA interference (RNAi) vector (*NbTRXh1*-RNAi) was constructed, and then the stable transgenic *N. benthamiana* plants downregulating *NbTRXh1* expression were generated. Intriguingly, compared with the non-transgenic plants, the *NbTRXh1*-RNAi transgenic plants displayed leaf mosaic developmental phenotype and the accumulation of endogenous NbTRXh1 protein was almost undetectable in the two transgenic RNAi lines 3E and 2F analyzed (Supplemental Figure S6, A and B). Then, BSMV was inoculated to *NbTRXh1*-RNAi transgenic and non-transgenic plants by agroinfiltration. The protein gel results showed that the accumulation of viral-encoded TGB1, CP, and γb proteins were increased in *NbTRXh1*-RNAi transgenic plants compared with those in non-transgenic plants (Figure 3E), suggesting that NbTRXh1 negatively regulates BSMV infection.

### NbTRXh1 specifically suppresses BSMV cell-to-cell movement

To determine whether the BSMV replication or cell-to-cell movement were impeded by NbTRXh1, we inoculated BSMV_γb-GFP_ (RNAα + RNAβ + RNAγ_γb-GFP_) or its movement-deficient mutant (RNAα + RNAγ_γb-GFP_, specifically initiate replication) into *N. benthamiana* plants which transiently overexpressed *NbTRXh1*-RNAi construct or empty vector (EV) using half-leaf method. The protein gel results showed that NbTRXh1 interferes with viral cell-to-cell movement instead of replication as evidenced by the similar accumulation of γb proteins in movement-deficient mutant virus-inoculated plants, whereas higher protein accumulation was detected in BSMV_γb-GFP_-inoculated *NbTRXh1*-RNAi plants (Figure 4A). Given that BSMV replicase αa recruits γb to chloroplasts replication sites for robust viral replication (Zhang et al., 2017; Jin et al., 2018), we also test the effect of NbTRXh1 on the interaction between γb and αa. The results showed that NbTRXh1 protein did not affect the stability of the γb-αa replication complex (Supplemental Figure S7), which is consistent with the lack of impact of NbTRXh1 on BSMV replication.

**Figure 4.**
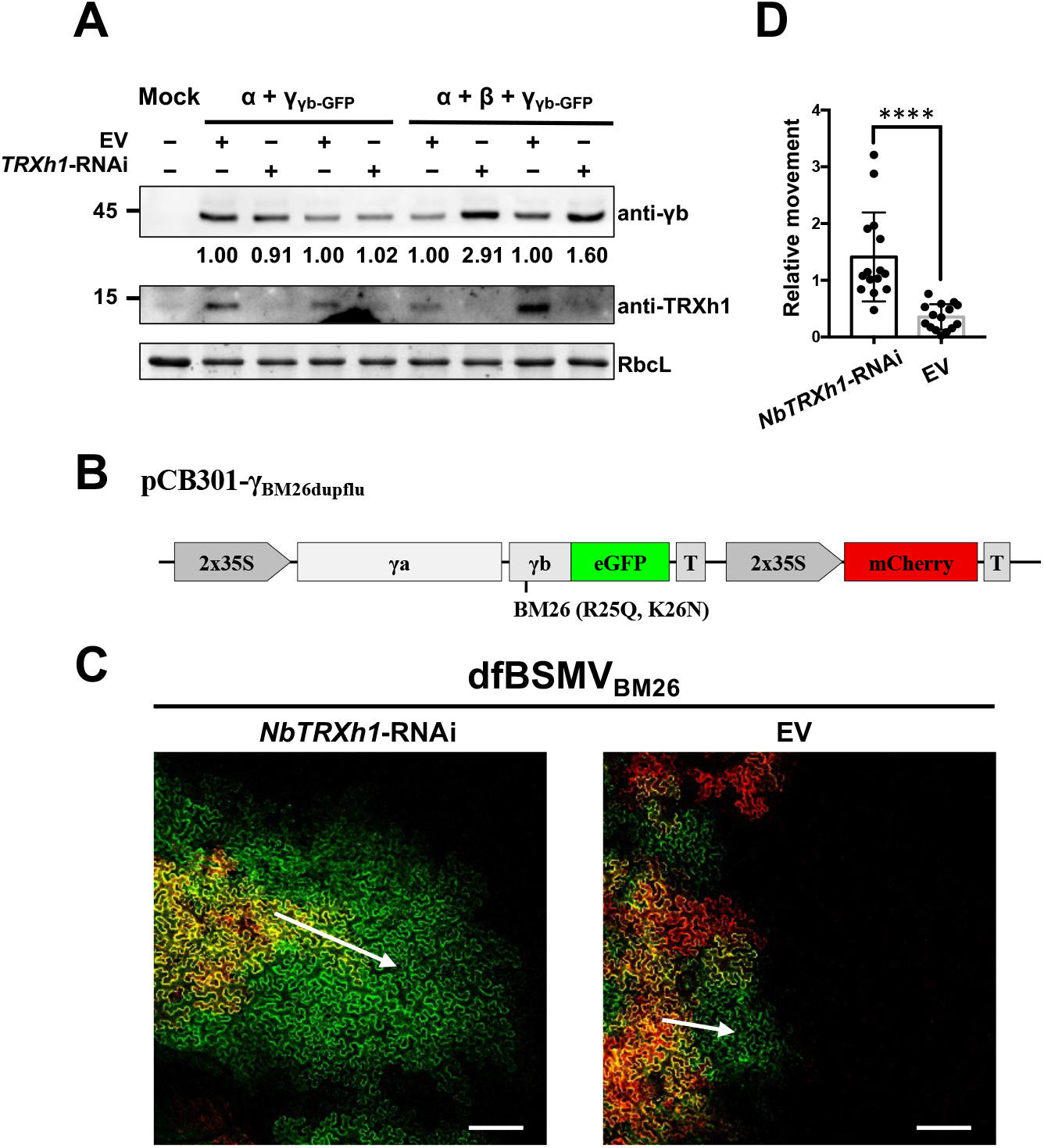
NbTRXh1 inhibits specifically BSMV cell-to-cell movement. **(A)** Western blot of BSMV accumulation in *NbTRXh1*-RNAi- or mock-inoculated *N. benthamiana* plants. *Agrobacterium* harboring the corresponding binary vectors were infiltrated into *N. benthamiana* plants by half-leaf method on the first day. BSMV RNAα and RNAγ_γb-GFP_ or BSMV RNAα, RNAβ, and RNAγ_γb-GFP_ were further inoculated into the previously agroinfiltrated leaves. Three days later, protein accumulation level of BSMV γb and endogenous NbTRXh1 was analyzed by using specific antibodies indicated on the right. RbcL was used as loading control. Experiments were repeated twice. **(B)** Schematic representation of pCB301-γ_BM26dupflu_ used for the dfBSMV_BM26_ reporter system. **(C)** Analysis of the effect of NbTRXh1 on BSMV movement. Images were captured at 2.5 dpi and the most representative images were displayed. The red signal shows primary agroinfiltrated area and the GFP signal outside the red region identifies secondary tissue invasion, which reflect the viral movement capacity. Bars, 200 μm. Arrows indicate the direction of viral cell-to-cell movement. **(D)** Quantification of BSMV movement shown in Figure 4C. The areas of green and red fluorescence were measured by ImageJ software. To quantify the relative movement ability, the GFP only area was divided by the corresponding red fluorescent area, and the results were analyzed by Student’s *t* test (n = 15, ****, *p* < 0.0001).

Next, we further used BSMV duplex fluorescence (dfBSMV) reporter system (Li et al., 2018; Jiang et al., 2020) to confirm the effect of NbTRXh1 on BSMV cell-to-cell movement. Since the viral suppressor of RNA silencing (VSR) activity of γb affects the silencing efficiency of hairpin RNA-mediated RNAi, a VSR-deficient dfBSMV_BM26_ mutant (pCB301-α + pCB301-β + pCB301-γ_BM26dupflu_, Figure 4B) was co-infiltrated with *NbTRXh1*-RNAi construct or EV into *N. benthamiana* by using half-leaf method. The results revealed that the viral cell-to-cell movement was significantly enhanced in *NbTRXh1*-silenced leaves compared with EV-infiltrated leaves (Figure 4C and 4D).

Altogether, these results indicate that NbTRXh1 through its interaction with γb targets specifically the viral cell-to-cell movement phase to inhibit BSMV infection.

### NbTRXh1 has no distinct effect on VSR activity of γb

BSMV γb is a well-characterized VSR that is of vital importance in viral pathogenesis (Donald and Jackson, 1994; Bragg and Jackson, 2004; Jiang et al., 2021). To test if NbTRXh1 impairs the BSMV infection by interfering with γb VSR activity, NbTRXh1 was co-expressed with γb and a positive-sense GFP (sGFP) (Zhang et al., 2018) in *N. benthamiana* leaves. At 3 dpi and 6 dpi, similar GFP fluorescence signal and GFP protein accumulation were observed in tissues expressing NbTRXh1 or EV (Supplemental Figure S8, A and B), demonstrating that NbTRXh1 does not affect VSR activity of γb protein.

A previous work has shown that NbTRXh2 (renamed as NbTRXh9.2 in Figure 1D) targets *Bamboo mosaic virus* (BaMV) TGB2 protein in *N. benthamiana*, which results in a restricted BaMV movement (Chen et al., 2018). Together with our results, we speculate that the molecular mechanisms of NbTRXh1-mediated antiviral defense might not via directly compromising the pro-viral functions of the γb protein *per se*, but rather a plant antiviral defense process.

### γb diminishes SA antiviral pathway via suppressing NbTRXh1 reductase activity

Since the reductase activity of NbTRXh1 is required for an antiviral strategy against BSMV (Figure 3B and 3D), which provide us with a hint that anti-BSMV role of NbTRXh1 might functionally linked with SA-mediated defense in *N. benthamiana*. To verify whether NbTRXh1 is involved in SA-mediated signaling pathway, six-leaf stage of *NbTRXh1*-RNAi transgenic or non-transgenic *N. benthamiana* plants were treated with 0.5 mM SA. The results showed that the transcript levels of *PR1* and *PR2* were significantly decreased in *NbTRXh1*-RNAi transgenic plants, indicating that NbTRXh1 is involved in SA-mediated defense pathway (Figure 5A and 5B).

**Figure 5.**
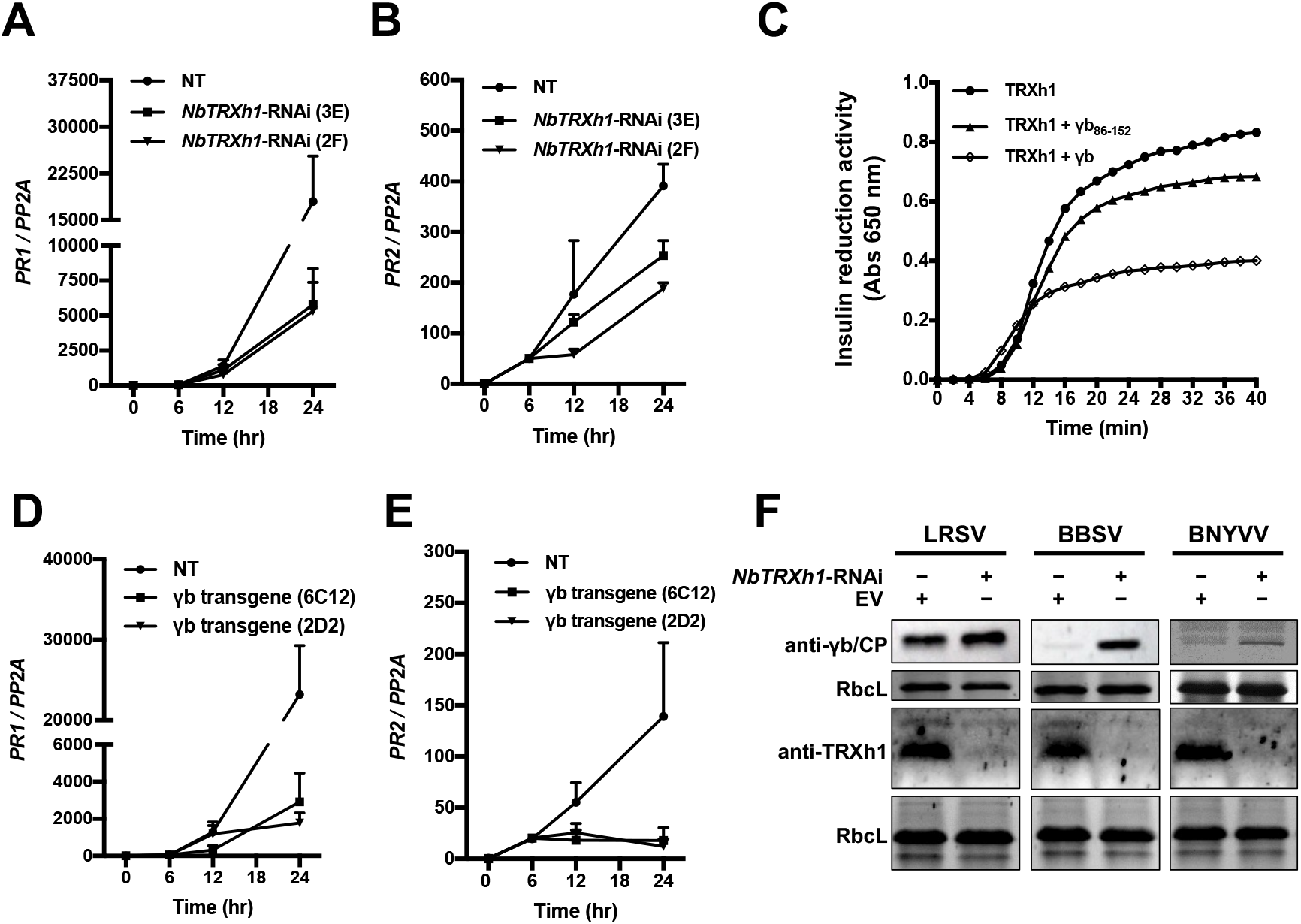
γb interferes with NbTRXh1-mediated defense. **(A-B)** NbTRXh1 is involved in SA defense pathway in *N. benthamiana* plants. Total RNA was extracted for RT-qPCR analysis of transcript levels of *PR1* and *PR2* in non-transgenic and *NbTRXh1*-RNAi transgenic (lines 3E and 2F) *N. benthamiana* plants after 0.5 mM SA treatment at 0 hpi, 6 hpi, 12 hpi, and 24 hpi, respectively. Values represent the average of three independent experiments, and error bars indicate Standard Error. **(C)** γb compromises NbTRXh1 reductase activity. 20 μM NbTRXh1 protein was used in this assay and insulin reduction activity was measured by reading absorbance at 650 nm. Experiments were performed thrice with similar tendency. **(D-E)** γb suppresses the expression of NPR1 downstream defense genes. Total RNA was extracted for RT-qPCR analysis of transcript levels of *PR1* and *PR2* in non-transgenic and γb-transgenic *N. benthamiana* plants (lines 6C12 and 2D2) upon 0.5 mM SA treatment at 0 hpi, 6 hpi, 12 hpi, and 24 hpi, respectively. Values represent the average of three independent experiments, and error bars indicate Standard Error. **(F)** Western blot analysis of LRSV γb, BBSV CP, and BNYVV CP proteins accumulation in *NbTRXh1*-silenced or mock-inoculated *N. benthamiana* plants at 3 dpi. RbcL was used as loading control. Experiments for each viral protein were repeated at least twice.

Next, we performed an *in vitro* insulin reduction assay (Holmgren, 1979) and the result showed that the recombinant NbTRXh1-His possesses reductase activity *in vitro* as evidenced by significantly increased absorbance values over time (Supplemental Figure S9). To verify whether γb subvert SA defense pathway by disturbing the enzymatic activity of NbTRXh1, the reductase activity of recombinant NbTRXh1-His were measured in the presence of GST-γb or GST-γb_86-152_ (a mutant that abolish NbTRXh1 binding capacity) proteins (Supplemental Figure 10). As expected, the result showed that the full-length γb protein, but not γb_86-152_, can suppress the NbTRXh1 reductase activity *in vitro* (Figure 5C).

Previous studies showed that the reductase activity of AtTRXh3 and AtTRXh5 orchestrate the *PR* gene expression to initiate plant defense in *Arabidopsis* (Kinkema et al., 2000; Tada et al., 2008). To further investigate whether γb affects the SA downstream defense genes expression, the γb-transgenic *N. benthamiana* plants (Yang et al., 2018) were treated with 0.5 mM SA and the transcript levels of *PR1* and *PR2* were analyzed at 0 h, 6 h, 12 h, 24 h after treatment. The RT-qPCR results showed that γb dramatically inhibited the SA downstream defense genes expression (Figure 5D and 5E).

Together, these data suggest that γb can compromise the SA downstream defense genes expression by directly inhibiting NbTRXh1 reductase activity.

### NbTRXh1-mediated defense responses against diverse plant viruses

To investigate whether NbTRXh1-mediated defense responses may be shared among members of highly diverged classes of plant viruses, we selected *Lychnis ringspot virus* (LRSV, genus *Hordeivirus*) (Jiang et al., 2018), *Beet black scorch virus* (BBSV, genus *Betanecrovirus*) (Wang et al., 2018), and *Beet necrotic yellow vein virus* (BNYVV, genus *Benyvirus*) (Jiang et al., 2019) to test the effect of NbTRXh1 on these viruses infections in *NbTRXh1*-RNAi construct- or EV-infiltrated *N. benthamiana* by half-leaf method. The results showed that the accumulation of these viruses increased in *NbTRXh1*-silenced half-leaves compared with those mock-inoculated one (Figure 5F). Intriguingly, previous studies has shown that the CaTRXh1-cicy suppresses the geminivirus infection in pepper (Luna-Rivero et al., 2016), the NtTRXh3 protein also negatively regulates TMV and *Cucumber mosaic virus* (CMV) infection in *N. tabacum* (Sun et al., 2010), the NbTRXh2 inhibits BaMV cell-to-cell movement in *N. benthamiana* (Chen et al., 2018); together with our results, we suggest that h-type TRXs have general roles to defend against both RNA and DNA virus infections in *planta*.

## Discussion

To successfully establish infections, pathogens have to battle with host defense system all the time. SA defense signaling pathway is one of the most important part of plant immunity network to defend against biotrophic and hemi-biotrophic pathogens (Ding and Ding, 2020). Most plants maintain low SA levels during normal growth and development, however, upon pathogen infection, SA is rapidly synthesized via two independent pathways through phenylalanine ammonia lyase (PAL) and isochorismate synthase (ICS); subsequently, NPR1, NPR3, and NPR4 perceive SA concentration changes and activate plant immunity (Peng et al., 2021). In this study, we found that BSMV infection activate SA defense pathway in *N. benthamiana*; exogenous SA treatment significantly inhibits BSMV accumulation (Figure 1). These results indicate that SA defense pathway is also involved in plant to defend against BSMV.

Upon recognition of pathogens, TRXs proteins sense the cytosolic redox status change and quickly switch the NPR1 oligomers-to-monomers (Tada et al., 2008); then, the monomers will be transported into nucleus and act as transcriptional coactivators to initiate downstream defense genes expression (Kinkema et al., 2000). Therefore, TRXhs-catalyzed NPR1 nuclear translocation process is an essential step in the SA defense pathway. Owing to the important role of TRXhs in plant immune system, several bacterial and fungal effectors have been identified to hijack TRXhs to evade plant defense system. For example, *Ralstonia solanacearum* employs an effector protein, RipAY, which exploits cytosolic TRXh proteins for self-activation and subsequent suppression of pathogen-associated molecular pattern (PAMP)-triggered immunity (PTI) (Mukaihara et al., 2016). A necrotrophic pathogen, *Cochliobolus victoriae*, confers host susceptibility by hijacking a TRXh5-dependent NB-LRR resistance pathway (Lorang et al., 2012). *Xanthomonas oryzae* pv. *oryzae* effector XopI disrupts NPR1-mediated resistance via proteasomal degradation of OsTRXh2 in rice (Ji et al., 2020). Likewise, in our study, we also found that viral encoded proteins could subvert TRXh-mediated plant defense. We provide direct evidences to indicate that NbTRXh1 inhibits BSMV infection through its reductase activity (Figure 3 and 4); more importantly, BSMV γb directly interacts with NbTRXh1 to weaken its reductase activity and optimize BSMV infection performance (Figure 2 and 5). Since NbNPR1 differs from AtNPR1 (Maier et al., 2011), whether NbTRXh1 is functionally linked to NbNPR1 and how NbTRXh1 regulates the SA signaling transduction need to be further investigated.

Replication and movement are two important steps on viral infection (Heinlein, 2015). To survival, plants have evolved versatile defense strategies to impair viral replication and/or movement phase. Several studies found that SA hormone has defense roles in both viral replication and movement stages. For example, SA treatment interefers with TMV RNA accumulation by cell type-specific effects on TMV replication and movement (Chivasa et al., 1997) (Murphy and Carr, 2002); in addition, SA inhibits the binding of cytosolic glyceraldehyde 3-phosphate dehydrogenase (GAPDH) protein to the negative RNA strand of *Tomato bushy stunt virus* (TBSV), thereby suppressing viral replication (Tian et al., 2015). In this study, we provide direct evidence to show that BSMV γb protein targets NbTRXh1 to block SA signaling transduction, which in turn specifically optimize viral cell-to-cell movement phase (Figure 4), indicating a negative role of SA in BSMV cell-to-cell movement. Since SA is a crucial molecular signal in the regulation of callose biosynthesis and deposition at plasmodesmata (PD) (Zavaliev et al., 2011; Wang et al., 2021) as well as pathogen-induced PD closure, it is possible that NbTRXh1 inhibits the BSMV intercellular movement via SA-mediated PD closure but this needs further investigation. Considering distinct movement patterns for the same virus in monocot and dicot plants (Lawrence and Jackson, 2001), future research should also include whether homologous TRXh1 proteins function in the same way to block BSMV cell-to-cell movement in monocot plants. Our results also demonstrate that NbTRXh1 reduces the accumulation of some other RNA viruses such as LRSV, BBSV, and BNYVV (Figure 5F). Although the specific infection process in which NbTRXh1 participates remains unclear, it suggests that TRXs could be a potential targets hijacked by different plant viruses. This study reveal that BSMV γb protein promotes viral movement in *N. benthamiana* via dampening the key plant SA defense pathway, however, whether other plant viruses also encode γb-like proteins to subvert this signaling pathways is still unclear.

Strikingly, we recently identified a NADPH-dependent thioredoxin reductase C (NTRC) involved in chloroplast antioxidant defense responses to disturb BSMV replication at chloroplasts replication sites (Wang et al., 2021). Intriguingly, the NTRC protein contains a thioredoxin domain (Trx-D) at its C-terminus, which also plays a negative role in BSMV replication when fused with a chloroplast transit peptide (cTP). Given that TRXs and Trx-D-containing proteins are also widely distributed in different organelles of the cell, such as chloroplasts, mitochondria, etc. (Mata-Pérez and Spoel, 2019; Kumari et al., 2021), and which usually are co-opted by different viruses at viral replication sites (Netherton and Wileman, 2011), it opens a question whether other TRXs or Trx-D containing proteins also have similar antiviral functions in plant defense system.

## Meterials and Methods

### Plasmid construction

The BSMV-related plasmids used in this study were described previously (Zhang et al., 2017; Hu et al., 2019; Jiang et al., 2020).

Full-length coding sequence (CDS) of *NbTRXh1* was amplified from cDNA of *N. benthamiana* and cloned into different expression vectors. Briefly, *NbTRXh1* was amplified by specific primers TRXh1-F and TRXh1-R, and inserted upon pET-30a plasmid digestion using same restriction enzymes for insulin reduction activity assay. For BiFC assay, the CDS of *TRXh1* was amplified with TRXh1-F2 and TRXh1-R2 containing *Bam*HI and *Xho*I sites and inserted into pSPYCE and pSPYNE digested with the same restriction enzymes. For co-IP assay, the CDS of TRXh1 was amplified with TRXh1-F3 and TRXh1-R3 containing *BamH*I and *Spe*I sites and inserted into pMDC32 to generate pMDC32-TRXh1-3xFlag. For pull-down assay, the full-length CDS of *TRXh1* were amplified by corresponding primers with *Bam*HI and *Xba*I sites and inserted into pMAL-c2x to generate prokaryotic expression vectors.

For TRXh1_SS_ mutant, site directed mutagenesis with TRXh1_SS_-F and TRXh1_SS_-R primers were used to generate *TRXh1_SS_* sequence and then inserted into different vectors (the same restriction enzymatic sites with TRXh1). For NPR1-3xFlag construct, full-length CDS of *NbNPR1* was amplified with NPR1-3xFlag-F and NPR1-3xFlag-R primers and then inserted into pMDC32-3xFlag plasmid at *Kpn*I and *Spe*I sites. To generate a plasmid construct that exogenously expresses hairpin RNAs targeting *NbTRXh1*, the gene-specific insert was constructed to transcribe a hairpin structure consisting of a 233 bp sequence from the target transcript with a spacer of 234 bp of random sequence for the hairpin loop, followed by the reverse complement of the original 233 bp. The fragment was integrated into pSK-In vector at *Kpn*I/*Sal*I and *Pst*I/*Spe*I sites, respectively, and then the fragments containing the forward and reverse NbTRXh1 sequences were amplified and inserted into pMDC32 vector at *Kpn*I and *Spe*I sites.

All the primers used for plasmid construction are listed in Supplemental Table S1, the correctness of all the plasmids were verified by DNA sequencing.

### Insulin reduction assay

The full-length CDS of NbTRXh1 was cloned into pET-30a to obtain purified NbTRXh1-his recombinant protein. The recombinant protein was used to perform insulin reduction assay as described previously with little modification (Holmgren, 1979). Briefly, different amounts of NbTRXh1 fusion proteins were added to the reaction buffer containing 25 mM Tris-HCl (pH 7.5), 1 mM EDTA, 200 μM bovine insulin and 0.5 mM DTT. The reduction activity was monitored at 650 nm at room temperature by using a spectrophotometer.

### Northern blotting

Northern blot analyses were performed as described previously (Zhang et al., 2017). Approximately 15 μg RNA and 4 μg RNA were loaded for endogenous *NbTRXh1* gene hybridization and viral RNA hybridization, respectively.

### Pull-down assay

Approximately 3 μg protein was incubated in binding buffer [25 mM Tris-HCl (pH 7.7), 150 mM NaCl, 0.2% (v/v) glycerol, 0.6% (v/v) TritonX-100, 1 x cocktail (Roche, Cat. # 11697498001)] containing 30 μl GST beads in a total volume of 600 μl. After 3 hours incubation at 4°C, beads were centrifugated and washed with washing buffer [20 mM Tris-HCl (pH 7.7), 250 mM NaCl, 0.2% (v/v) glycerol, 0.6% (v/v) TritonX-100] at 4°C for six times. Then SDS loading buffer was added and the beads were boiled for 10 min before subjected to western blotting.

### Agroinfiltration

All binary vectors used were transformed into *Agrobacterium tumefaciens* strain EHA105 as described previously (Cao et al., 2015). In transient expression assays, suspensions of *Agrobacterium* carrying NbTRXh1 or its mutant constructs were infiltrated at OD_600_ = 0.5. For hairpin RNA-mediated RNA interference approaches, the *Agrobacterium* harboring the *NbTRXh1*-RNAi construct was infiltrated at OD_600_ = 0.5. For Co-IP, *Agrobacterium* containing each construct were infiltrated at OD_600_ = 0.4. For BSMV and LRSV inoculation, the OD_600_ of *Agrobacterium* containing each RNA segment is 0.3. For BBSV inoculation, the OD_600_ of the inoculum was 0.3; and for BNYVV RNA1, RNA2, and RNA4 is 0.05 for each construct.

### Co-immunoprecipitation assays

Co-IP assays were performed as described previously (Jiang et al., 2022). Briefly, plant tissue expressing different proteins was collected at 3 dpi and extracted in extraction buffer [25mM Tris-HCl (pH 7.5), 150mM NaCl, 1mM EDTA, 10% (v/v) glycerol, 2% (w/v) polyvinylpolypyrrolidone, 1% protease inhibitor Cocktail (Roche, Cat. # 11697498001), 10mM dithiothreitol]. The crude extracts were centrifuged at 8000 *g* for 30 minutes and the supernatants were collected and incubated with FLAG beads (Sigma). After 4 hours incubation at 4°C, the beads were collected by centrifugation at 800 *g* for 1 min and washed six times with IP buffer [25 mM Tris-HCl, 150mM NaCl, 10% (v/v) glycerol, 0.1% (v/v) Tween 20] at 4°C during 10 min per wash. The pelleted beads were boiled for 10 min and analyzed by western blotting.

### Quantitative reverse transcription PCR (RT-qPCR)

RT-qPCR analyses were performed as described previously (Jiang et al., 2020). Notably, both oligo(dT)_20_ and specific PR2-R primers were added into the reaction buffer when generating cDNA. Primers used for RT-qPCR analyses were listed in Supplemental Table 2.

## Accession numbers

Sequence data from this work can be found in updated *N. benthamiana* database (https://ora.ox.ac.uk/objects/uuid:f34c90af-9a2a-4279-a6d2-09cbdcb323a2) (Kourelis et al., 2019) or the *Arabidopsis* Information Resource (www.Arabidopsis.org) under the following accession numbers: NbTRXh1 (NbD046153.1), NbTRXh1.2 (NbD034443.1), NbTRXh1.3 (NbD029648.1), NbTRXh2.1 (NbD052469.1), NbTRXh2.2 (NbD015627.1), NbTRXh2.3 (NbE03054613.1), NbTRXh2.4 (NbD015628.1), NbTRXh9.1 (NbD027853.1), NbTRXh9.2 (NbD031089.1), AtTRXh1 (AT3G51030), AtTRXh2 (AT5G39950), AtTRXh3 (AT5G42980), AtTRXh4 (AT1G19730), AtTRXh5 (AT1G45145), AtTRXh7 (AT1G59730), AtTRXh8 (AT1G69880), AtTRXh9 (AT3G08710), and AtTRXh10 (AT3G56420).

## Supplemental Information

The following materials are available in the online version of this article.

**Supplemental Figure S1. Viral accumulation of BSMV, BSMV_BM26_, and BSMV_Δγb_ at 3 dpi.**

Actin was used as loading control. Antibodies are indicated on the right.

**Supplemental Figure S2. Sequence alignment of TRXhs from *N. benthamiana* and *Arabidopsis*.**

The conserved reductase domain was indicated with blue line.

**Supplemental Figure S3. Western blot to confirm the protein accumulation in BiFC assay.**

Arrowheads indicate the fusion proteins. Anti-GFP antibody is indicated on the right.

**Supplemental Figure S4. BSMV γb protein cannot affect the subcellular localization of NbTRXh1 during viral infection.**

**(A)** Different combinations were agroinfiltrated into *N. benthamiana* leaves and the images were captured at 2.5 dpi. Bars, 20 μm. Dashed squares indicate the position of zoom-in panels. **(B)** Protein accumulation from samples in Figure S4A. RbcL was used as loading control. Antibodies are indicated on the right. Numbers on the left indicate molecular weight in kDa.

**Supplemental Figure S5. GST pull-down assay to analyze the *in vitro* γb_BM26_ and NbTRXh1 interaction.**

Purified GST-γb_BM26_ or GST proteins were incubated with NbTRXh1-His. After being immunoprecipitated with glutathione-Sepharose beads, proteins were detected by western blot with anti-His and anti-GST antibodies. Antibodies are indicated on the right of the corresponding blot.

**Supplemental Figure S6. Developmental phenotype and molecular analysis of *NbTRXh1*-RNAi transgenic lines.**

**(A)** Pictures were taken at 4 weeks. Bars, 5 cm. **(B)** NbTRXh1 protein accumulation in *NbTRXh1* RNAi lines. Total protein was extracted and analyzed by western blot. Arrowhead indicates the endogenous NbTRXh1 protein. RbcL was used as loading control. Mr indicates protein molecular marker.

**Supplemental Figure S7. The effect of NbTRXh1 on the γb-αa interaction.**

Different constructs shown on the top of the figure were coinfiltrated into *N. benthamiana* leaves. At 3 dpi, total protein was extracted and incubated with anti-Flag beads. Input and IP products were analyzed by western blot with anti-GFP or anti-Flag antibodies. Antibodies are indicated on the right of the corresponding blot. Numbers on the left indicate molecular weight in kDa.

**Supplemental Figure S8. NbTRXh1 impacts on the VSR activity of BSMV γb.**

**(A)** *N. benthamiana* leaves coexpressing BSMV γb protein and sGFP together with EV or untagged NbTRXh1. Images were taken under long-wave UV light at 3 dpi or 6 dpi. **(B)** Analysis of GFP protein accumulation in Figure S8A. Anti-GFP, anti-γb, and anti-TRXh1 antibodies were used to detect the corresponding protein accumulation. RbcL was used as loading control.

**Supplemental Figure S9. *In vitro* analysis of reductase activity of the recombinant NbTRXh1 protein.**

The NbTRXh1-His protein was purified from *E. coli* and different amounts of protein (5 μM, 10 μM, and 20 μM) were used to test its reductase activity. Absorbance values at 650 nm were measured during 60 min.

**Supplemental Figure S10.** Input proteins used for insulin reductase assay in Figure 5C.

Arrowheads indicate the fusion proteins.

**Supplemental Table S1.** Primers used in this study.

## Acknowledgments

We would like to thank Dr. Laura Medina-Puche (Centre for Plant Molecular Biology, Eberhard Karls University, Germany) for critical reading and thorough editing of this manuscript. We thank the members of Li lab for their helpful discussions and suggestions. No conflict of interest declared.

